# Intestinal epithelial MHC class II regulation by HDAC3 instructs microbiota-specific CD4+ T cells

**DOI:** 10.1101/2022.01.01.473735

**Authors:** Emily M. Eshleman, Tzu-Yu Shao, Vivienne Woo, Taylor Rice, Jordan Whitt, Laura Engleman, Sing Sing Way, Theresa Alenghat

## Abstract

Aberrant immune responses to resident microbes promote inflammatory bowel disease and other chronic inflammatory conditions. However, how microbiota-specific immunity is controlled in mucosal tissues remains poorly understood. Here, we find that mice lacking epithelial expression of microbiota-sensitive histone deacetylase 3 (HDAC3) exhibit increased accumulation of commensal-specific CD4^+^ T cells in the intestine, provoking the hypothesis that epithelial HDAC3 may instruct local microbiota-specific immunity. Consistent with this, microbiota-specific CD4^+^ T cells and epithelial HDAC3 expression were concurrently induced following early-life microbiota colonization. Further, epithelial-intrinsic ablation of HDAC3 promoted T cell driven-colitis and primed development of pathogenic commensal-specific Th17 cells. Mechanistically, HDAC3 was essential for MHC class II (MHCII) expression by the intestinal epithelium, and epithelial-intrinsic MHCII actively limited commensal-specific Th17 cells and prevented microbiota-triggered inflammation. Remarkably, HDAC3 enabled the microbiota to induce MHCII on epithelial cells and limit the number of commensal-specific T cells in the intestine. Collectively, these data reveal a central role for an epithelial histone deacetylase in controlling development of tissue-intrinsic T cells that recognize commensal microbes and drive pathologic inflammation.

## Introduction

The gastrointestinal tract is home to trillions of microorganisms, collectively termed the microbiota, which form symbiotic relationships with mammalian cells and play a significant role in mediating health and disease. Extensive experiments utilizing broad-spectrum antibiotics or germ-free (GF) mouse models have revealed the necessity for the microbiota in the development and function of the host immune system (1–4). Microbiota interactions are particularly impactful during early-life, as critical immune education and calibration occur within the first few years of life (5–8). Indeed, disturbances or perturbations in microbiota composition or colonization during this early-life window have been associated with long-lasting changes in immunity that can predispose to the development of chronic inflammatory disorders including asthma, allergy, and inflammatory bowel diseases (IBD) (1, 4, 7).

Despite the symbiotic nature of the host-microbiota relationship, the close association and abundance of microbes at mucosal surfaces pose a potential risk to stimulate pathologic inflammation. This requires that intestinal immune responses must be tightly regulated to allow protective immunity against invading pathogens, while limiting inflammatory responses towards innocuous commensal microbes. Commensal bacteria drive regulatory T cell differentiation (9), and microbiota-reactive effector and memory T cells are present in both mice and humans (10–14). These commensal-specific T cells promote barrier function by inducing protective cytokines and providing cross-reactivity to pathogens (13, 15, 16). However, aberrant immune responses to the microbiota also trigger inflammatory conditions, such as IBD (1, 17–19). In mouse models of colitis, intestinal microbes drive inflammation, in part, by stimulating microbiota-reactive CD4^+^ T cells (20–22). Further, microbiota-specific CD4^+^ T cells in IBD patients have been shown to be functionally altered and produce more pro-inflammatory cytokines such as IL-17, compared to T cells from healthy patients (13, 23–27). However, the mechanisms instructing tissue-intrinsic regulation of commensal-specific CD4^+^ T cells remains poorly understood.

Intestinal epithelial cells (IECs) reside at the direct interface between the microbiota and immune cells, and are thus uniquely poised to instruct local immunity in response to microbial antigens. IECs express pathogen recognition receptors that sense microbial signals and regulate intestinal immune responses via secretion of cytokines, chemokines, and growth factors (1, 28). While specialized microfold cells and goblet cell-associated antigen passages in the epithelium deliver antigens to underlying antigen presenting myeloid cells (29–31), IECs are also equipped with classical antigen processing and presentation pathways that can regulate immune responses (32, 33). However, there remains limited understanding of IEC-directed mechanisms that coordinate healthy microbiota-immune relationships.

Beyond canonical sensors, epithelial expression of the epigenetic-modifying enzyme histone deacetylase 3 (HDAC3) has recently been found to respond to the microbiota and regulate mammalian metabolism, intestinal homeostasis, and inflammation (34–37). Here, we found that microbiota-dependent expansion of commensal-specific CD4^+^ T cells following initial microbiota colonization occurred concurrently with epithelial expression of HDAC3. Given this relationship, we examined whether IEC-intrinsic HDAC3 regulates antigen specificity directed to the microbiota. Indeed, loss of HDAC3 expression in IECs resulted in intestinal inflammation and accumulation of microbiota-specific Th17 cells that were regulated by epithelial MHCII. Interestingly, germ-free analyses demonstrated that HDAC3 was necessary for microbiota to induce epithelial MHCII-dependent regulation of microbiota-specific T cells. Taken together, these data reveal that the microbiota induce commensal tolerance and limit inflammation by directing epithelial control of tissue-intrinsic T cells.

## Results

### Epithelial HDAC3 expression limits commensal-specific CD4^+^ T cells in the intestine

Microbiota colonization begins at birth with the complexity and density of the microbiota increasing most significantly during infancy (38–40). Microbiota exposure early in life is required for effective immune education and disruption during this window can increase chronic inflammatory disorders (39–43). While it has been predicted that initial colonization at birth must direct development of local T cell specificity to commensal microbes, this has not been directly tested. Therefore, to investigate whether early-life microbiota colonization itself instructs commensal-specific T cell responses in the intestine, CD4^+^ T cells from the large intestine of germ-free (GF) and conventionally-housed (CNV) neonatal pups were first analyzed for specificity to microbial antigens using MHCII-tetramers loaded with commensal-flagella peptide, cBir1. Within the first week of life, GF and CNV pups exhibited a similarly low abundance of commensal-specific cBir1^+^ CD4^+^ T cells in the large intestine (**Figure 1, A and B**). However, microbiota colonization increases by weaning (41) and, interestingly, 3-week old pups reared in the presence of microbes already demonstrated an accumulation of cBir1^+^ CD4^+^ T cells compared to age-matched GF controls (**Figure 1, A and B**). Thus, as predicted, early microbiota-derived signals are essential for initial expansion of intestinal microbiota-specific CD4^+^ T cells.

**Figure 1.**
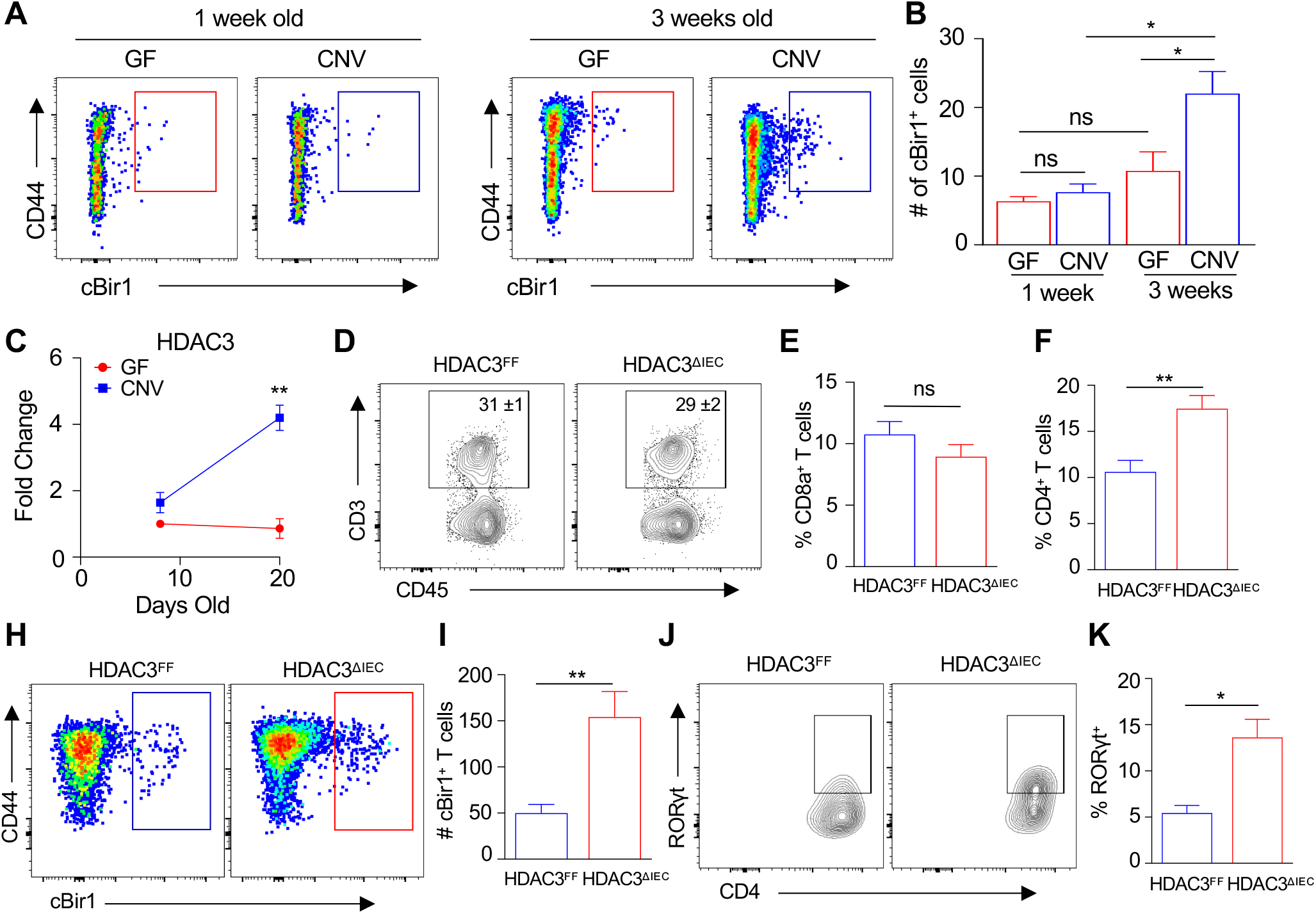
Epithelial HDAC3 expression limits commensal-specific CD4^+^ T cells in the intestine. (**A, B**) Number of cBir1^+^ tetramer-specific CD4^+^ T cells isolated from large intestine of neonatal GF and CNV pups. (**C**) mRNA expression of HDAC3 in IECs isolated from large intestine of GF and CNV pups. (**D**) Frequency of total intestinal CD3^+^. (**E**) Frequency of total intestinal CD8a^+^ and (**F**) CD4^+^ T cells. (**H, I**) Number of cBir1^+^ tetramer-specific CD4^+^ T cells and (**J, K**) frequency of RORγt^+^ cBir1^+^ CD4^+^ T cells in large intestine of HDAC3^FF^ and HDAC3^ΔIEC^ mice. cBir1 ^+^ tetramer cells are gated on live, CD45^+^, lineage (CD11b, B220, Ly6G, CD11c, CD8a)^-^, CD4^+^ cells. Data represent at least two independent experiments, 3-4 mice per group. *p<0.05, **p<0.01, ns=not significant.

Dysregulated commensal-specific T cells accumulate in patients with intestinal inflammation (13), suggesting that impairment in pathways that regulate these T cell subsets can impact susceptibility to pathologic inflammation. Interestingly, IECs from IBD patients express decreased levels of the HDAC3 enzyme (34) and, consistent with findings in other facilities (34), mice lacking IEC-intrinsic expression of HDAC3 (HDAC3^ΔIEC^ mice) displayed increased susceptibility to chronic intestinal inflammation (34) characterized by rectal prolapse (**Supplemental Figure 1A**), increased levels of the inflammatory biomarker, lipocalin-2 (**Supplemental Figure 1B**), infiltration of inflammatory cells (**Supplemental Figure 1, C-E**), and pathology consistent with chronic inflammation (**Supplemental Figure 1F**). Strikingly, epithelial expression of HDAC3 was also dramatically induced in 3-week old pups relative to younger 1-week old pups (**Figure 1C**). However, age-matched pups raised under GF conditions demonstrated low epithelial HDAC3 expression that was unaltered with age (**Figure 1C**), indicating that microbiota colonization increases early-life epithelial HDAC3 expression in the intestine.

The temporal link between epithelial-intrinsic HDAC3 expression and expansion of commensal-specific CD4^+^ T cells following microbiota colonization provoked the hypothesis that IEC-intrinsic HDAC3 may regulate microbiota-specific T cell immunity. In the intestinal lamina propria, total CD3^+^ T cells were unaltered in HDAC3^ΔIEC^ mice compared to wildtype floxed littermate controls (HDAC3^FF^) (**Figure 1D**). Interestingly, though, loss of epithelial HDAC3 resulted in elevated intestinal CD4^+^ T cells, but not CD8a^+^ cells, (**Figure 1, E and F**) that exhibited increased cBir1^+^ commensal-specificity (**Figure 1, H and I**). These cBir1^+^ CD4^+^ T cells were characterized by a higher frequency of pro-inflammatory RORγt^+^ Th17 cells in HDAC3^ΔIEC^ mice compared to floxed controls (**Figure 1, J and K**). Taken together, these data indicate that epithelial HDAC3 plays a critical role in regulating intestinal microbiota-specific Th17 cells.

### CD4^+^ T cells from mice lacking epithelial HDAC3 induce severe colitis

To determine whether dysregulated CD4^+^ T cells in HDAC3^ΔIEC^ mice promoted intestinal inflammation, the T cell transfer model of chronic colitis was employed in which purified CD4^+^ T cells were isolated from HDAC3^FF^ and HDAC3^ΔIEC^ mice and transferred into Rag1^-/-^ mice (**Figure 2A**). In contrast to Rag1^-/-^ recipients who received CD4^+^ T cells from control mice, Rag1^-/-^ recipients receiving CD4^+^ T cells from HDAC3^ΔIEC^ mice displayed more significant weight loss (**Figure 2B**), colonic shortening (**Figure 2C**), and severe colitis pathology characterized by inflammatory cell infiltration, crypt hyperplasia, and mural thickening (**Figure 2, D and E**). Furthermore, luminal concentration of the inflammatory biomarker, lipocalin-2, was also significantly upregulated in Rag1^-/-^ mice that received CD4^+^ T cells from HDAC3^ΔIEC^ mice (**Figure 2F**). Therefore, CD4^+^ T cells from mice that specifically lack IEC-intrinsic HDAC3 are sufficient to induce intestinal inflammation in genetically-susceptible hosts.

**Figure 2.**
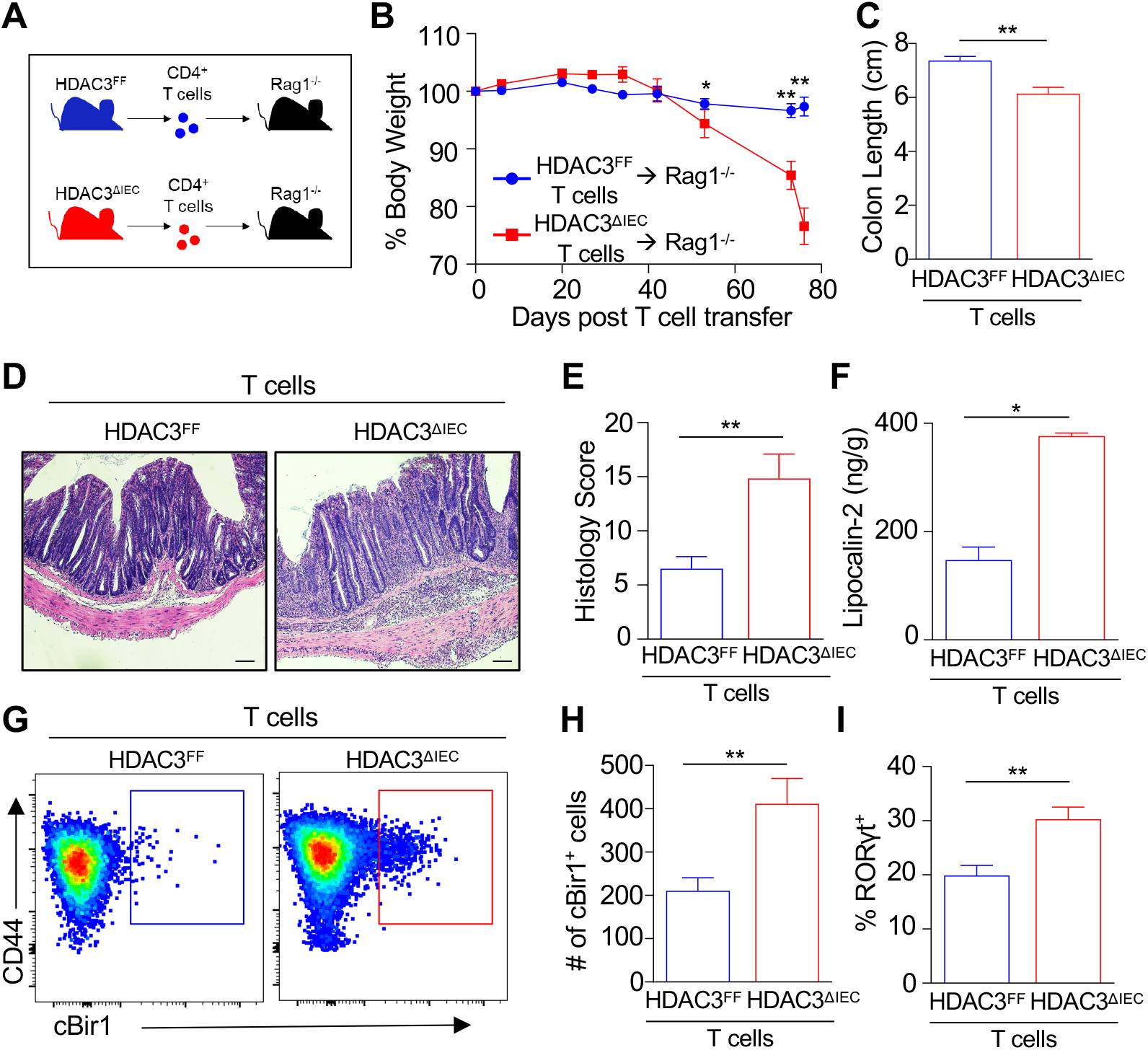
CD4^+^ T cells from mice lacking epithelial HDAC3 induce severe colitis. (**A**) Experimental schematic of T cell colitis model with naïve CD4^+^ T cells isolated from HDAC3^FF^ and HDAC3^ΔIEC^ mice transferred into Rag1^-/-^ hosts. (**B**) Change in body weight, (**C**) colon length, (**D**) H&E-stained colonic sections of Rag1^-/-^ hosts that received T cells from HDAC3^FF^ or HDAC3^ΔIEC^ mice, scale bars 20 μM. (**E**) Histological scoring of sections in (D). (**F**) Fecal concentration of lipocalin-2. (**G, H**) Number of cBir1 ^+^ tetramer-specific CD4^+^ T cells in large intestine of Rag1^-/-^ hosts that received T cells from HDAC3^FF^ or HDAC3^ΔIEC^ mice. Gated on live, CD45^+^, lineage (CD11b, B220, Ly6G, CD11c, CD8a)^-^, CD4^+^. (**I**) Frequency of RORγt^+^ cBir1^+^ CD4^+^ T cells. Data represent at least two independent experiments, 3-4 mice per group. *p<0.05, **p<0.01.

To next investigate whether colitogenic T cells isolated from HDAC3^ΔIEC^ mice were responding to commensal antigens, intestinal microbiota-reactive CD4^+^ T cells were evaluated. Rag1^-/-^ recipients that received CD4^+^ T cells from HDAC3^ΔIEC^ mice exhibited elevated microbiota-specific cBir1 ^+^ CD4^+^ T cells compared to hosts that received HDAC3^FF^ cells (**Figure 2, G and H**). In addition, a higher frequency of commensal-specific T cells differentiated into inflammatory RORγt^+^ Th17 cells when they originated from HDAC3^ΔIEC^ mice compared to HDAC3^FF^ littermate controls (**Figure 2I**). Collectively, these data indicate that epithelial HDAC3 expression is required for regulation of CD4^+^ T cell development in the intestine, as loss of epithelial HDAC3 resulted in increased microbiota-specific colitogenic CD4^+^ T cells.

### HDAC3 regulates surface expression of MHC class II on intestinal epithelial cells

Antigen presentation via MHCII is critical for instructing antigen-specific CD4^+^ T cell responses. However, the frequency of total MHCII^+^ intestinal hemopoietic cells (**Figure 3A**) and classical APCs, including MHCII^hi^ intestinal dendritic cells and macrophages, were unaffected by epithelial HDAC3 deletion (**Figure 3, B and C**). Surprisingly, though, we found that EpCAM^+^ IECs, and not CD45^+^ hematopoietic cells, expressed the majority of MHCII at the intestinal-microbiota interface (**Figure 3, D and E**). IECs respond to microbial cues, so we next assessed whether the microbiota plays a role in regulating IEC-intrinsic MHCII expression. Consistent with other studies (44–48), IECs isolated from CNV mice displayed higher surface expression of MHCII compared to GF mice (**Figure 3, F and G**). Further, neonatal GF pups maintained low epithelial expression of H2-Ab1, the gene which encodes the beta-chain of MHCII (**Figure 3H**). However, epithelial H2-Ab1 expression was robustly induced following initial microbiota colonization (**Figure 3H**), similar to regulation of HDAC3 (Figure 1C). These data indicate the necessity of the microbiota in controlling IEC-intrinsic MHCII and suggest that HDAC3 may regulate epithelial MHCII. Therefore, surface expression of MHCII protein was compared on IECs isolated from the large intestine HDAC3^FF^ and HDAC3^ΔIEC^ mice. Interestingly, loss of HDAC3 expression dramatically reduced MHCII expression on IECs (**Figure 3, I and J**). These findings reveal a critical regulatory role for HDAC3 in directing surface expression of MHCII on intestinal epithelial cells.

**Figure 3.**
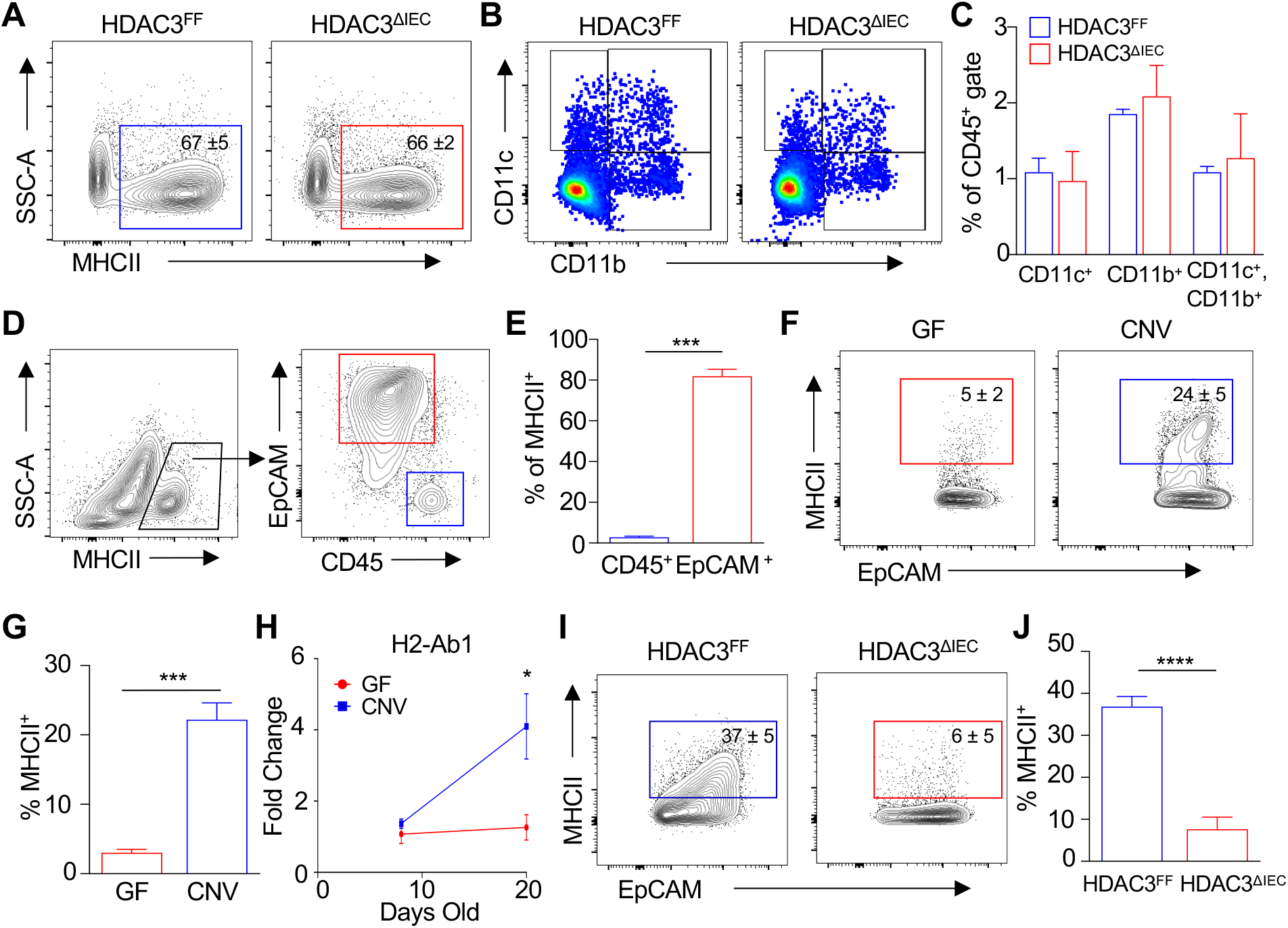
HDAC3 regulates surface expression of MHC class II on intestinal epithelial cells. (**A**) Frequency of total MHCII^+^ cells in colon lamina propria. (**B, C**) Frequency of dendritic cells and macrophages in large intestinal lamina propria of HDAC3^FF^ and HDAC3^ΔIEC^ mice. Gated on live, CD45^+^, MHCII^+^. (**D**) MHCII^+^ expressing cells at colonic luminal surface. (**E**) Frequency of total MHCII^+^ cells in (D). (**F, G**) Frequency of MHCII^+^ EpCAM^+^ cells in large intestine of GF and CNV mice. (**H**) mRNA expression of H2-Ab1 in IECs isolated from large intestine of GF and CNV pups. (**I, J**) Frequency of MHCII^+^ EpCAM^+^ cells in large intestine of HDAC3^FF^ and HDAC3^ΔIEC^ mice. Data represent at least three independent experiments, 3-4 mice per group. *p<0.05, ***p<0.001, ****p<0.0001.

### Epithelial MHCII limits commensal-specific CD4^+^ T cells and intestinal inflammation

While epithelial MHCII has been suggested to function in both protective or detrimental immune responses (44, 48–50), its role in regulating commensal-specific T cell responses and microbiota-triggered inflammation has not been directly examined. Therefore, to test this, we generated an IEC-specific MHCII knockout model (MHCII^ΔIEC^) by crossing floxed H2-Ab1 mice (MHCII^FF^) to mice expressing Cre recombinase downstream of the villin promoter (51, 52). Significant reduction in MHCII expression in IECs of MHCII^ΔIEC^ mice was confirmed by mRNA (**Supplemental Figure 2A**) and surface protein analyses (**Supplemental Figure 2B**) and, importantly, loss of MHCII expression did not occur on CD45^+^ cells in MHCII^ΔIEC^ mice (**Supplemental Figure 2C**). In contrast to the activating role of MHCII on classical APCs, loss of IEC-intrinsic MHCII led to elevated accumulation of commensal-specific cBir1 ^+^ CD4^+^ T cells in the colon of MHCII^ΔIEC^ mice relative to floxed control mice (**Figure 4, A and B**). In order to examine whether this commensal-specific response occurred with another microbial antigen, mice were also colonized with *Candida albicans* expressing 2W1S_55-68_ variant of I-Ea epitope (**Figure 4C**) (53–55). Consistent with regulation of cBir1 ^+^ CD4^+^ T cells, MHCII^ΔIEC^ mice displayed increased numbers of *C. albicans*-2W1S-specific CD4^+^ T cells relative to MHCII^FF^ mice (**Figure 4, D and E**) with similar levels of *C. albicans* colonization. Collectively, these data reveal that IEC-intrinsic MHCII does in fact control commensal-specific CD4^+^ T cell levels in the intestine.

**Figure 4.**
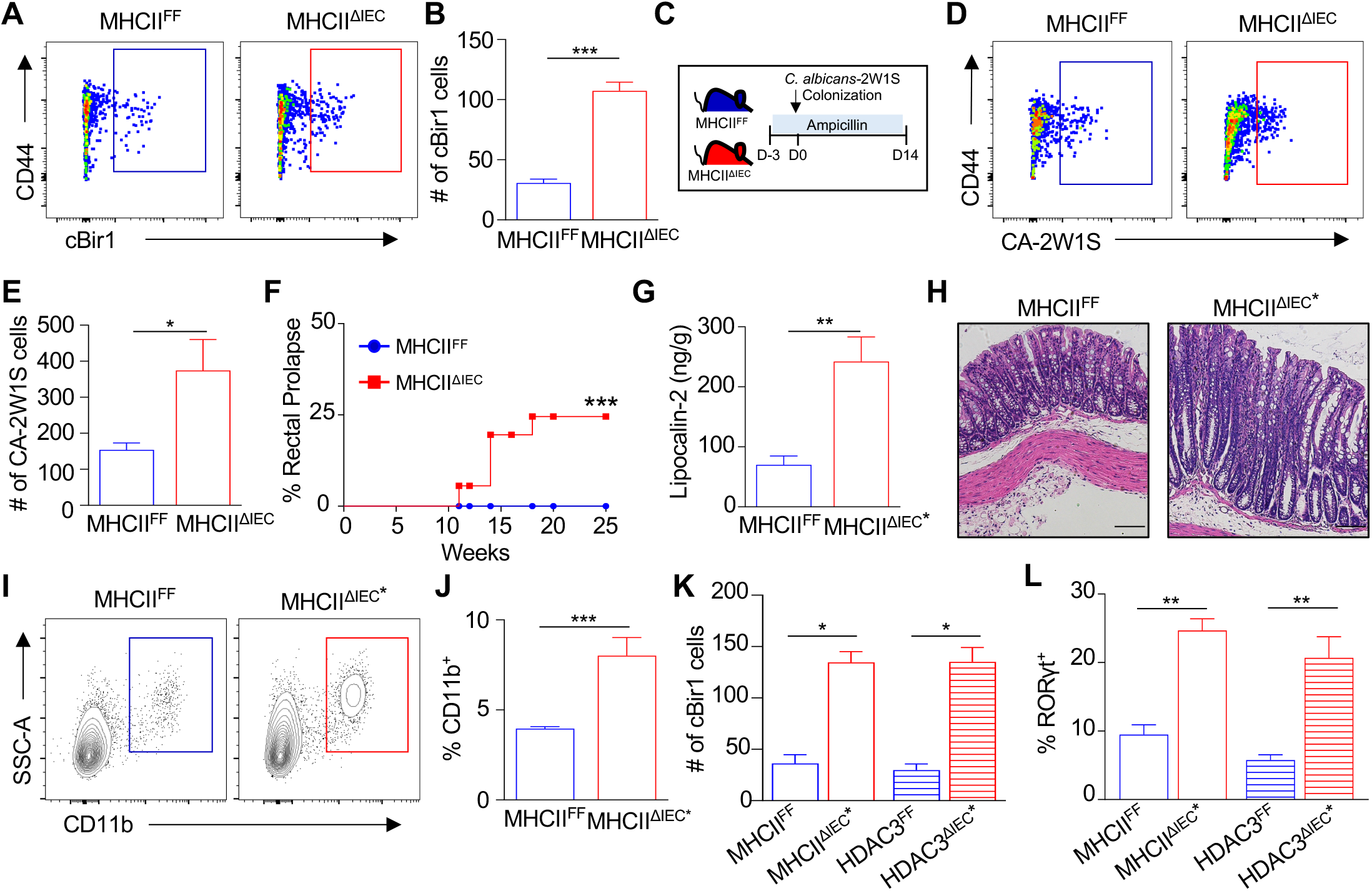
Epithelial MHCII limits commensal-specific CD4^+^ T cells and intestinal inflammation. (**A, B**) Number of cBir1^+^-specific CD4^+^ T cells in large intestine of MHCII^FF^ and MHCII^ΔIEC^ mice. (**C**) Diagram of *2W1S-Candida albicans* commensal colonization. (**D, E**) Number of *C. albicans*-2W1S^+^-specific CD4^+^ cells in large intestine of MHCII^FF^ and MHCII^ΔIEC^ mice. (**F**) Frequency of rectal prolapse in MHCII^FF^ and MHCII^ΔIEC^ mice. (**G**) Lipocalin-2 levels in stool of MHCII^FF^ and prolapsed MHCII^ΔIEC^ mice (MHCII^ΔIEC^*). (**H**) H&E-stained colonic sections of MHCII^FF^ and MHCII^ΔIEC^* mice, scale bars 20 μM. (**I, J**) Frequency of myeloid cell infiltrate in MHCII^FF^ and MHCII^ΔIEC^* mice. (**K**) cBir1^+^-specific CD4^+^ T cells and (**L**) frequency of RORγt^+^ cBir1^+^-specific CD4^+^ T cells in large intestine of control and prolapsed MHCII^ΔIEC^ and HDAC3^ΔIEC^ mice. Gated on live, CD45^+^, lineage (CD11b, B220, Ly6G, CD11c, CD8a)^-^, CD4^+^. Data are representative of at least two independent experiments, 3-6 mice per group. *p<0.05, **p<0.01, ***p<0.001.

Mice lacking IEC-intrinsic HDAC3 exhibited increased susceptibility to chronic intestinal inflammation (Figure S1). Remarkably, similar to HDAC3^ΔIEC^ mice, mice lacking epithelial MHCII also displayed an increased incidence of rectal prolapse with age (**Figure 4F**), indicative of intestinal inflammation. Furthermore, prolapsed MHCII^ΔIEC^ mice (MHCII^ΔIEC^*) demonstrated increased fecal lipocalin-2 (**Figure 4G**), pathology consistent with chronic colitis (**Figure 4H**), and increased infiltration of myeloid cells (**Figure 4, I and J**). Interestingly, MHCII^ΔIEC^ mice and HDAC3^ΔIEC^ mice with an increased propensity to prolapse were characterized by elevated levels of commensal-specific CD4^+^ T cells (**Figure 4K**) that reflected commensal-specific RORγt^+^ inflammatory Th17 cells (**Figure 4L**). Collectively, these data suggest that epithelial HDAC3 may be critical in regulating microbiota-induced MHCII-directed commensal-specific immunity and inflammation.

### HDAC3 enables microbiota to regulate epithelial dependent commensal-specific immunity

To test the requirement for the microbiota in MHCII-dependent intestinal inflammation and commensal-specific immunity, MHCII^FF^ and MHCII^ΔIEC^ mice were raised in the absence and presence of broad-spectrum antibiotics that significantly deplete commensal bacteria (56). Consistent with earlier results (Figure 4F), MHCII^ΔIEC^ mice exhibited an increased prevalence of rectal prolapse (**Figure 5A**), whereas depletion of the microbiota prevented spontaneous intestinal inflammation in MHCII^ΔIEC^ mice (**Figure 5A**). Furthermore, microbiota-replete MHCII^ΔIEC^ mice exhibited accumulation of commensal-specific Th17 cells (**Figure 5B**) and increased levels of IL-17 (**Figure 5C**). However, antibiotic-treated MHCII^ΔIEC^ mice did not demonstrate increased commensal-specific Th17 cells (**Figure 5B**) or IL-17 expression (**Figure 5C**) relative to MHCII^FF^ mice. Taken together, these data show that the microbiota is necessary in triggering epithelial MHCII-dependent intestinal inflammation and commensal-specific immune responses.

**Figure 5.**
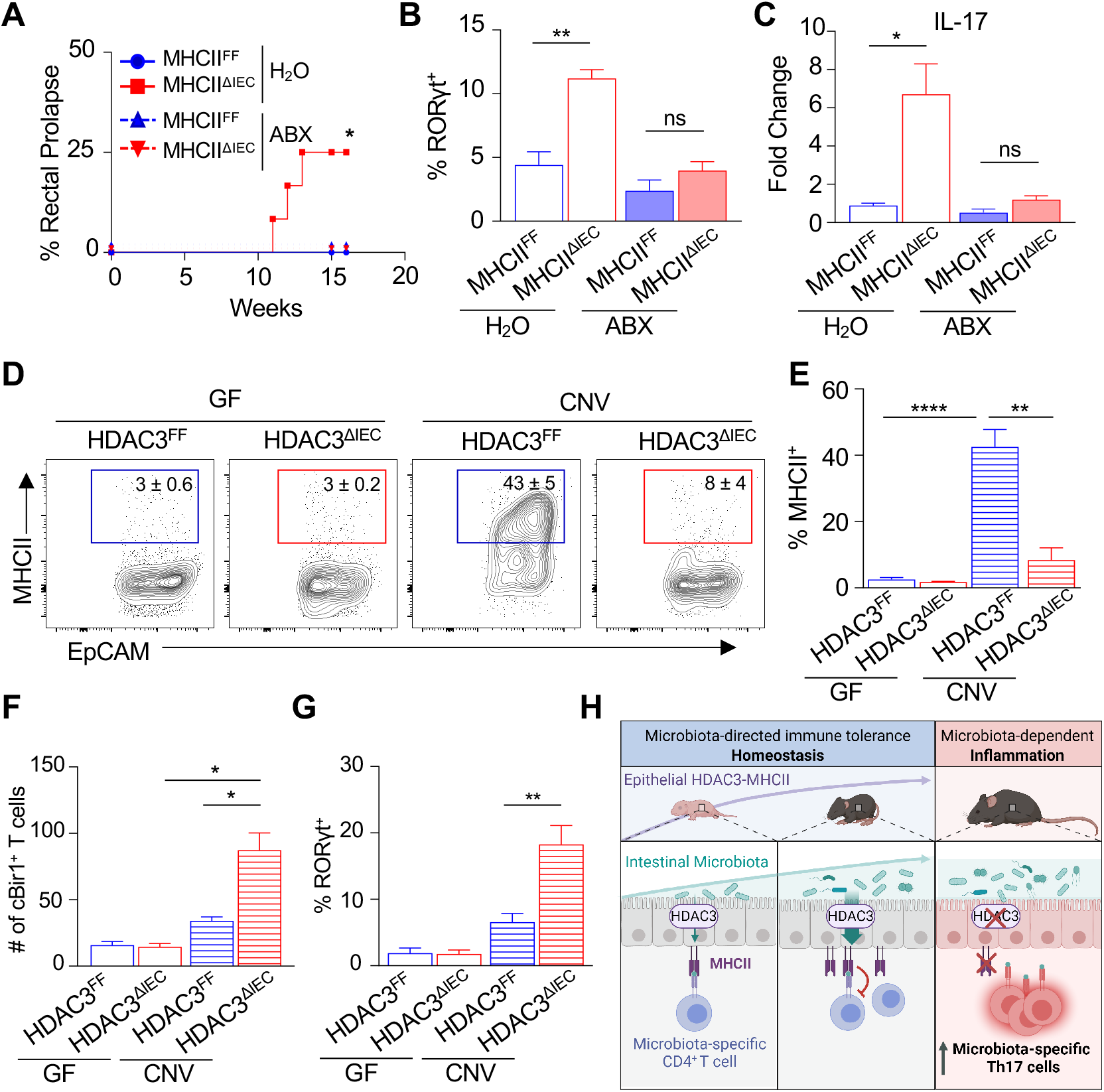
HDAC3 enables microbiota to regulate epithelial dependent commensal-specific immunity. (**A**) Frequency of rectal prolapse in control and antibiotic (ABX)-treated MHCII^FF^ and MHCII^ΔIEC^ mice. (**B**) Frequency of RORγt^+^ cBir1^+^-specific CD4^+^ T cells and (**C**) IL-17 mRNA expression in large intestine of control and ABX-treated MHCII^FF^ and MHCII^ΔIEC^ mice. (**D, E**) Frequency of MHCII^+^ EpCAM^+^ cells in large intestine of GF- and CNV-HDAC3^FF^ and HDAC3^ΔIEC^ mice. (**F**) Number of cBir1^+^ tetramer-specific CD4^+^ T cells and (**G**) frequency of RORγt^+^ cBir1 ^+^ T cells isolated from GF- and CNV-HDAC3^FF^ and HDAC3^ΔIEC^ mice. cBir1^+^ tetramer cells are gated on live, CD45^+^, lineage (CD11b, B220, Ly6G, CD11c, CD8a)^-^, CD4^+^. (**H**) Microbiota drive selftolerance early in life through upregulation of an epithelial HDAC3 dependent-MHCII pathway that limits commensal-specific Th17 cell accumulation and pathologic inflammation. Data are representative of or pooled from at least two independent experiments, 3-6 mice per group. *p<0.05, **p<0.01, ****p<0.0001, ns=not significant.

Therefore, to test whether HDAC3 is necessary in mediating this regulation of epithelial and immune cells by the microbiota, GF-derived HDAC3^FF^ and HDAC3^ΔIEC^ mice were compared to CNV-HDAC3^FF^ and HDAC3^ΔIEC^ mice. Consistent with earlier data employing wildtype CNV and GF mice (Figure 3F, G), IEC-intrinsic MHCII expression was significantly induced in CNV-HDAC3^FF^ controls compared to GF-HDAC3^FF^ mice (**Figure 5, D and E**), whereas CNV-HDAC3^ΔIEC^ mice failed to upregulate MHCII (**Figure 5, D and E**). In GF mice, loss of epithelial HDAC3 expression had no impact on protein levels of MHCII on IECs (**Figure 5, D and E**), demonstrating the specific necessity for HDAC3 in mediating microbiota-dependent regulation of epithelial MHCII. Furthermore, CNV-HDAC3^ΔIEC^ mice exhibited elevated cBir1^+^ commensal-specific Th17 cells compared to CNV-HDAC3^FF^ littermate controls (**Figure 5, F and G**). However, GF-HDAC3^ΔIEC^ had similar levels of commensal-specific Th17 cells as GF-HDAC3^FF^ mice (**Figure 5, F and G)**. These data indicate the necessity for HDAC3 in enabling the microbiota to regulate commensal-specific immunity. Collectively, these findings reveal that microbiota colonization can induce commensal self-tolerance through an epithelial HDAC3-MHCII regulatory pathway that dampens commensal-specific immune responses directly in the local tissue environment (**Figure 5H**).

## Discussion

The microbiota is essential for the development and education of the host immune system. However, inappropriate immune reactions to the microbiota underlie several chronic inflammatory conditions, highlighting the necessity to understand how microbiota-specific immunity is controlled. In this study, we found that initial microbiota colonization led to upregulation of epithelial HDAC3 expression and expansion of commensal-specific T cells. Selective deletion of HDAC3 in IECs resulted in decreased epithelial MHCII expression and impaired regulation of commensal-specific CD4^+^ T cells that are controlled by epithelial MHCII. Importantly, these commensal-specific T cells developed into Th17 cells that promote microbiota-triggered pathologic inflammation. Furthermore, HDAC3 was necessary for microbiota to be able to induce epithelial MHCII-dependent regulation of microbiota-specific T cells. Therefore, the microbiota can direct self-tolerance by inducing a HDAC3-dependent epithelial MHCII pathway that regulates local commensal-specific T cells and susceptibility to microbiota-sensitive disease (**Figure 5H**).

Microbiota-specific CD4^+^ T cells are generated centrally in the thymus (57) and in peripheral mucosal tissues (14, 58–60). While auto-reactive CD4^+^ T cell are well known to be limited by thymic negative selection (61–63), the mechanisms controlling commensal-specific cells remain less well understood. Previous work described that negative selection of activated commensal-specific T cells can be mediated by MHCII-expressing group 3 innate lymphoid cells (ILC3) (64, 65). While ILC3s are relatively rare cells at the luminal surface of the intestine, we show that IECs are a major source of intestinal MHCII. Indeed, HDAC3-dependent MHCII expression specifically by epithelial cells limited accumulation of commensal-specific Th17 and thus, may be a dominant mechanism at the luminal surface for controlling commensal-specific CD4^+^ T cells and dampening pathogenic responses to the microbiota. Given the role for MHCII in ILC3s (64, 65), complementary and essential roles for antigen presentation pathways in distinct non-classical cells seem to be crucial for establishing and maintaining healthy intestine-intrinsic tissue homeostasis.

Loss of HDAC3-dependent epithelial MHCII led to elevated commensal-specific Th17 activity and increased susceptibility to microbiota-driven intestinal inflammation (**Figure 5H**). Patients with active IBD have functionally distinct microbiota-reactive CD4^+^ T cells, compared to those found in healthy controls, and secrete higher levels of IL-17 (13, 23, 25–27). IL-17 has been linked to development and exacerbation of several autoimmune and inflammatory conditions, including IBD (25, 27, 66). Furthermore, mouse models with disrupted IL-17 signaling are protected from intestinal inflammation (67, 68), supporting a pathogenic role for IL-17 in intestinal inflammation. However, clinical trials depleting IL-17 in IBD patients have been ineffective, and in some cases worsened disease (69). This pleotropic role for IL-17 in regulating intestinal homeostasis suggests that targeting specific IL-17 producers, instead of broadly neutralizing IL-17 itself, may be a more effective treatment. In fact, work has shown that therapeutics which target commensal-specific Th17 cells, while retaining IL-17 production from other cell types, reduced intestinal inflammation (70). Our data suggests that promoting epithelial MHCII expression through enhanced HDAC3 activity may further restrict pro-inflammatory commensal-specific T cells and promote healthy intestinal homeostasis.

Crosstalk between the microbiota and mammalian immune cells involves communication via pattern recognition receptor engagement and microbiota-derived metabolite signaling. Our data reveal a distinct level of epithelial regulation in which the microbiota-sensitive enzyme, HDAC3, regulates epithelial MHCII expression in the large intestine. Indeed, loss of epithelial HDAC3 was sufficient to abrogate microbiota-dependent MHCII expression in the intestine. Previous studies have linked the requirement for the microbiota in regulating IEC-intrinsic MHCII expression (44–48). Many of these studies have focused on how immune cells can induce epithelial MHCII through IFNγ signaling (44, 71, 72). In addition, disruption to canonical pattern recognition pathways leads to loss of intestinal barrier integrity and increased inflammation (73–75), and the toll-like receptor signaling adaptors MyD88/TRAF have been shown to induce epithelial MHCII expression, particularly in the small intestine (44). However, whether crosstalk between microbiota-sensing pathways and/or IFNγ signaling are required for HDAC3 regulation of epithelial MHCII and commensal-specific immunity remains to be determined.

HDAC3 is a multifaceted enzyme that can deacetylate histone and non-histone targets to alter gene expression and cellular functions. HDAC3 can be regulated by endogenous factors, dietary components, and microbiota-derived signals (34, 35, 76–78). While HDAC3 classically suppresses gene expression of direct targets via histone deacetylation, more recent work has shown that HDAC3 may also function as a transcriptional co-activator to induce gene expression through deacetylase-independent activity (79–84). Given that H2-Ab1 expression is decreased in HDAC3^ΔIEC^ mice, loss of HDAC3 may either directly increase expression of a repressor of H2-Ab1 or alter MHCII expression through a secondary or non-histone target.

Increasing evidence suggests that several tissue-specific, non-hematopoietic cells, including lung epithelial cells, skin keratinocytes, and fibroblasts express MHCII and the necessary machinery to regulate tissue-intrinsic CD4^+^ T cell responses (85–87). Given the ubiquitous nature of HDAC3, it is thus plausible that HDAC3 promotes MHCII-dependent pathways in other non-classical antigen presenting cells as well. Interestingly, tissue-specific deletion of HDAC3 has been associated with chronic inflammatory disease models for diabetes, heart disease, Alzheimer’s, and IBD (34, 35, 88–90). Importantly, our findings here uncover a fundamental new tenet for immune regulation whereby HDAC3 induction of non-hematopoietic epithelial MHCII is largely induced by the microbiota to dampen local self-directed immune responses. These findings reveal a central host mechanism that is utilized by the microbiota to instruct commensal-directed immunity, and suggest that this regulatory pathway can be targeted to treat chronic inflammatory conditions.

## Methods

### Mice

Conventionally-housed C57Bl/6J mice were purchased from Jackson Laboratories and maintained in our specific-pathogen free colony at Cincinnati Children’s Hospital Medical Center (CCHMC). Germ-free (GF) mice were maintained in flexible isolators in the CCHMC Gnotobiotic Mouse Facility, fed autoclaved feed and water, and monitored for absence of microbes. HDAC3^FF^ and HDAC3^ΔIEC^ were generated as described (34). H2-Ab1(MHCII)^FF^ were purchased from Jackson and maintained at CCHMC. MHCII^FF^ mice were crossed to C57BL/6J-Villin-Cre to generate MHCII^ΔIEC^ mice. Sex- and age-matched littermate controls were used for all studies. Mice were housed up to 4-per cage in a ventilated cage system in a 12-h light/dark cycle, with free access to water and food. For T cell transfer colitis model, 5 x 10^5^ naïve CD4^+^ T cells were isolated from the spleen and mesenteric lymph nodes of HDAC3^FF^ and HDAC3^ΔIEC^ mice via the MojoSort Mouse CD4 Naïve T Cell Isolation (BioLegend). Cell purity was confirmed by flow cytometry and T cells injected IP into age- and sex-matched Rag1^-/-^ recipients. *Candida albicans* colonization was conducted as previously described (53, 54). Briefly, mice were pre-treated with ampicillin-water (1-mg/mL) two days prior to colonization and maintained on ampicillin-water for the duration of the experiment. Mice were gavaged with recombinant *C. albicans* expressing 2W1S55-68 peptide (55). Mice were monitored for colonization by plating fecal CFUs and were harvested 14-days post-colonization. For antibiotic treatment, MHCII^FF^ and MHCII^ΔIEC^ mice pups were provided water supplemented with 1-mg/mL colistin (Sigma), 1-mg/mL ampicillin (Sigma) and 5-mg/mL streptomycin (Sigma) at weaning and maintained on antibiotics for 16-weeks. Antibiotic-water was refreshed every 7-10 days. All mouse studies were conducted with approval by Animal Care and Use Committees at CCHMC. These protocols follow standards enacted by the United States Public Health Services and Department of Agriculture. All experiments followed standards set forth by Animal Research: Reporting of In Vivo Experiments (ARRIVE).

### Cell Isolation

The large intestine was harvested, opened, and washed in PBS. For IECs, tissue was placed in pre-warmed strip buffer (PBS, 5% FBS, 1mM EDTA, 1mM DTT) and incubated at 37°C at 45-degree angle with shaking at 180-rpm for 15-min. For lamina propria isolation, tissue was washed with PBS to remove EDTA and DTT then incubated in pre-warmed digestion buffer (RPMI with 1-mg/mL Collagenase/Dispase (Sigma)) at 37°C at 45-degree angle with shaking at 180-rpm for 30-min. After incubation, the tissue was vortexed and passed through a 70*μ*M cell strainer.

### Flow cytometry

Cells were stained for flow cytometry using the following antibodies diluted in FACS Buffer (2% FBS, 0.01% Sodium Azide, PBS): BV711-anti-CD326 (EpCAM) (Clone:G8.8, BioLegend), BUV395-anti-CD45.2 (Clone:104, BD Biosciences), APC or FITC-anti-MHCII (Clone:M5/114.15.2, eBioscience), APC-eFluor-780-anti-CD4 (Clone:RM4-5, eBioscience), Pe-Cy7-anti-CD44 (Clone:IM7, eBioscience), AlexaFluor-647-anti-RORγt (Clone:Q31-378, BD Biosciences), PerCP-eFluor710 or APC-anti-CD8a (Clone:53-6.7, eBioscience), PerCP-Cy5.5-anti-CD3 (Clone: 17A2, eBioscience), PerCP-eFluor710-anti-B220 (Clone:RA3-6B2, eBioscience), PerCP-eFluor710-anti-Ly6G (Clone:1A8-Ly6g, eBioscience), PerCP-Cy5.5 or PE-anti-CD11b (Clone:M1/70, eBioscience), PerCP-Cy5.5-anti-CD11c (Clone:N418, eBioscience). Dead cells were excluded with the Fixable Aqua Dead Cell Stain Kit (Invitrogen). The BD Fix/Perm kit was used for intracellular staining. Class-II restricted tetramers (cBir1 and 2W1S) were PE-conjugated and were generated and provided by the NIH tetramer core. Samples were acquired on the BD LSR-Fortessa and analyzed with FlowJo Software (Treestar).

### RNA and qPCR analysis

RNA was isolated using the RNeasy Kit (Qiagen). For RNA from whole tissue, samples were homogenized in TRIzol. Chloroform was added for phase separation and RNA was precipitated by mixing with isopropanol. cDNA was synthesized using the Verso reverse transcriptase kit (Thermo Fisher) following the manufacturer’s protocol. Real-time PCR was performed using SYBR green (Applied Biosystems) and analyzed using the following murine primers sequences. HPRT forward 5’-GATTAGCGATGAACCAGGT-3’, HPRT reverse 5’-CCTCCCATCTCCTTCATGACA-3’, H2-Ab1 forward 5’-TGTGAGTCCTGGTGACTGCCATTA-3’, H2-Ab1 reverse 5’-TCGCCCATGAACTGGTACACGAAA-3’, IL-17 forward 5’-ACCGCAATGAAGACCCTGAT-3’, IL-17 reverse 5’-TCCCTCCGCATTGACACA-3’, HDAC3 forward 5’-TTGGTATCCTGGAGCTGCTT-3’, HDAC3 Reverse 5’-GACCCGGTCAGTGAGGTAGA-3’.

### Lipocalin-2 ELISA

Fecal pellets were homogenized in PBS at a concentration of 100-mg/mL then centrifuged at high-speed for 10-mins. Supernatants were collected and lipocalin-2 levels were determined using mouse Lipocalin-2/NGAL (R&D Systems) following the manufacturer’s instructions.

### Histological Analysis

Colonic tissue sections were fixed in 4% paraformaldehyde overnight at 4°C then paraffin embedded, sectioned, and stained with hematoxylin and eosin. For histological scoring, slides were evaluated based on the following parameters: immune cell infiltration (1-5), mucosal thickening/edema (1-5), crypt length (1-5), and crypt abscess/erosion (1-5).

### Statistical analysis

Results are expressed as mean ± SEM. Statistical significance was determined with the student’s t-test or one-way ANOVA with all data meeting the assumptions of the statistical test used. Results were considered significant at *p < 0.05; **p < 0.01; ***p < 0.001, ****p < 0.0001. Statistical significance was calculated using Prism version 7.0 (GraphPad Software).

## Author Contributions

Conceptualization, E.M.E., S.S.W., T.A.; Methodology, E.M.E, T.-Y.S., V.W., L.E., J.W., S.S.W., T.A.; Investigation, E.M.E., J.W., T.-Y.S., V.W., T.R.; Formal Analysis, E.M.E, T.-Y.S., T.R.; Writing, Review and Editing, E.M.E., S.S.W., T.A.

## Acknowledgements

We thank the Way, Qualls, and Deshmukh labs for useful discussions and members of the Alenghat lab for critical reading of the manuscript. We thank CCHMC Veterinary Services, Research Flow Cytometry Core, Confocal Imaging Core, and Pathology Research Core for services and technical assistance. This research is supported by the National Institutes of Health (R01DK114123, R01DK116868 to T.A., DP1AI131080 to S.S.W, and F32AI147591 to E.M.E.), and a Kenneth Rainin Foundation award to T.A. S.S.W. and T.A. each hold Investigators in the Pathogenesis of Infectious Disease Award from the Burroughs Wellcome Fund. This project is supported in part by PHS grant P30DK078392.

