## SupplementalFigures for "Intestinal epithelial MHC class II regulation by HDAC3 instructs microbiota-specific CD4+ T cells"

### Supplemental Figure 1

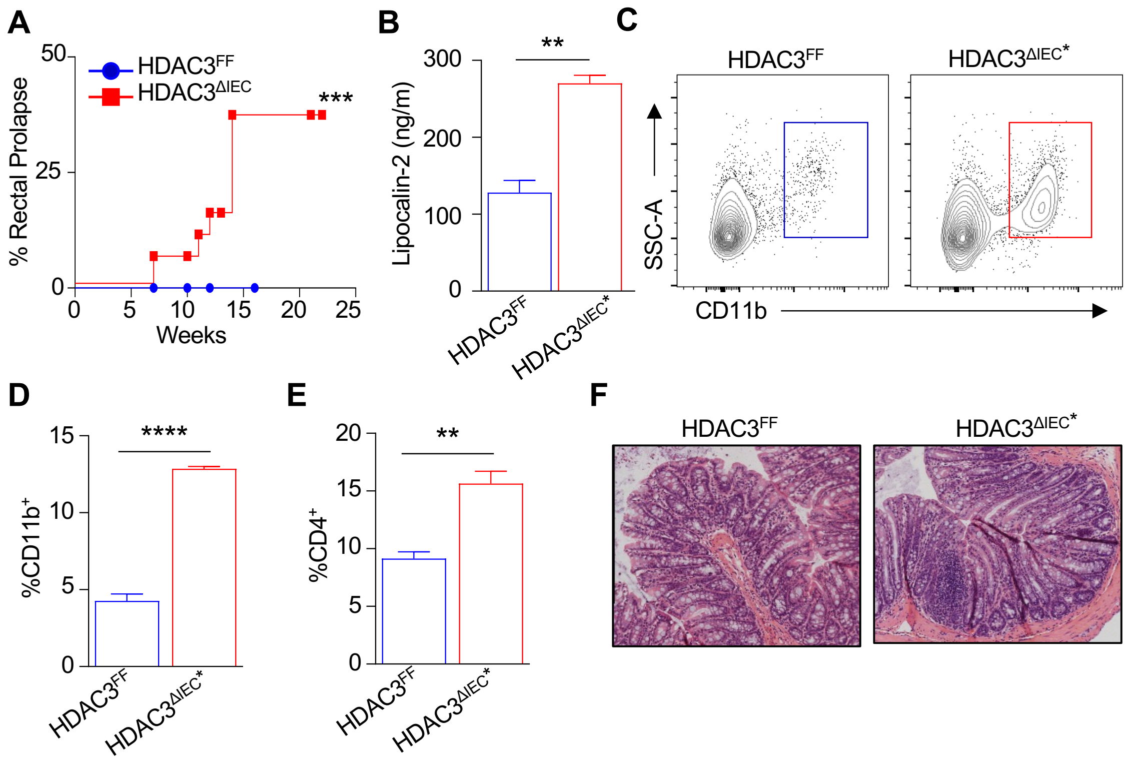


**Supplemental Fig. 1. Loss of epithelial HDAC3 expression increases spontaneous intestinal inflammation**. (**A**) Frequency of rectal prolapse in HDAC3$^{FF}$ and HDAC3$^{\Delta IEC}$ mice. (**B**) Fecal concentrations of lipocalin-2. (**C, D**) Frequency of CD11b$^{+}$ myeloid cells (Gated Live, CD45$^{+}$, CD3$^{-}$) and (**E**) frequency of intestinal CD4$^{+}$ T cells (Gated Live, CD45$^{+}$, CD3$^{+}$) in the large intestine of HDAC3$^{FF}$ and prolapse (*) HDAC3$^{\Delta IEC}$ mice. (**F**) H&E-stained colonic sections. Data represent at least three independent experiments, 3-4 mice per group. **$p<0.01$, ***$p<0.001$, ****$p<0.0001$.

**Supplemental Figure 2**

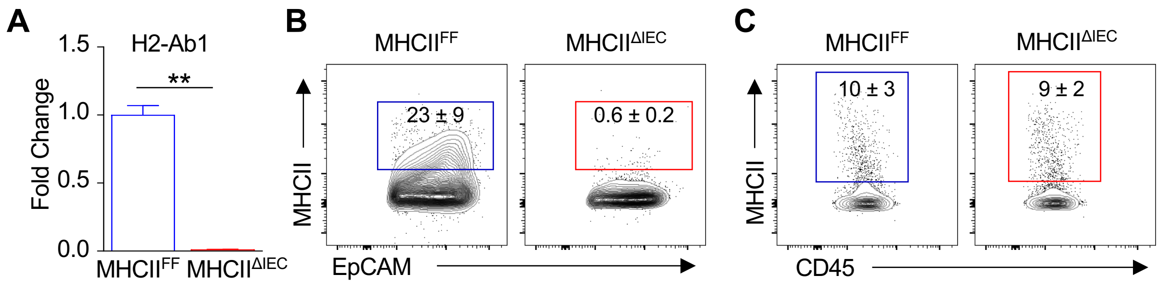


**Supplemental Fig. 2. MHCII$^{\Delta IEC}$ mice exhibit specific loss of MHCII on intestinal epithelial cells.** (**A**) H2-Ab1 mRNA expression in IECs isolated from the large intestine of MHCII$^{FF}$ and MHCII$^{\Delta IEC}$ mice. (**B**) Surface MHCII expression in EpCAM$^{+}$, CD45$^{-}$ cells from MHCII$^{FF}$ and MHCII$^{\Delta IEC}$ mice. (**C**) Surface MHCII expression in EpCAM$^{-}$, CD45$^{+}$ cells from MHCII$^{FF}$ and MHCII$^{\Delta IEC}$ mice. Data represent at least two independent experiments, 3-4 mice per group. $**p<0.01$.
